# Delineating JunB’s Crucial Function in Mature Th17 Cells through Inducible Targeted Protein Degradation

**DOI:** 10.1101/2025.01.06.631425

**Authors:** Charles Whitaker, Daiki Sasaki, Mio Miyagi, Masato Hirota, Miho Tamai, Ke Wang, Hsiao-Chiao Chien, Shukla Sarkar, Takeshi Toma, Naoyuki Taira, Hiroki Ishikawa

## Abstract

The AP-1 transcription factor JunB is essential for the differentiation of pathogenic T helper 17 (Th17) cells, which are key mediators of autoimmune diseases such as multiple sclerosis and colitis. While the importance of JunB during Th17 polarization is known, its role in mature Th17 cells—critical therapeutic targets in these diseases—remains unclear. In this study, we employed the dTAG system, a targeted protein degradation approach, to deplete JunB in Th17 cells generated both *in vitro* and *in vivo*. During pathogenic Th17 cell differentiation, JunB degradation replicated known effects of JunB deficiency, including reduced expression of interleukin (IL)-17A and the genes encoding RORγt (*Rorc*) and the IL-23 receptor (*Il23r*). In contrast, in mature pathogenic Th17 cells, JunB degradation downregulated *Il23r* without affecting IL-17A or *Rorc* expression. Furthermore, JunB degradation compromised the viability of mature pathogenic Th17 cells. Transcriptomic analyses revealed that JunB regulates distinct gene sets during Th17 polarization compared to mature Th17 cells. The gene *Inhba*, which encodes activin A, was identified as a JunB target in both stages. Supplementation with activin A restored IL-17A and *Rorc* expression during pathogenic Th17 cell differentiation. These findings demonstrate that JunB maintains mature pathogenic Th17 cell phenotypes, including IL-23 receptor expression, and supports pathogenic Th17 cell survival. As IL-23 signaling is crucial for sustaining pathogenic Th17 cells, targeting JunB may offer a therapeutic strategy to limit Th17-driven autoimmune inflammation.

## Introduction

T helper (Th) 17 cells are a subset of CD4+ T cells that play a critical role in host defense, particularly in the clearance of extracellular bacterial and fungal infections (1,2). However, Th17 cells are also major drivers of inflammation in autoimmune diseases such as multiple sclerosis, psoriasis, and inflammatory bowel disease (IBD) (3–5). Th17 cell differentiation is initiated by IL-6-mediated activation of STAT3, which drives the expression of effector cytokines, the master transcription factor retinoid-related orphan receptor γt (RORγt, encoded by *Rorc*), and the IL-23 receptor (IL-23R) (6,7). RORγt promotes the expression of key Th17 molecules, including IL-17A, IL-17F, and IL-23R, in both mouse and human cells (8,9). The upregulation of IL-23R allows differentiating Th17 cells to respond to IL-23 signals, which further activate STAT3, creating a positive feedback loop that supports Th17 cell maturation and maintenance (10).

Th17 cells can be classified into pathogenic (pTh17) and non-pathogenic (npTh17) subsets, depending on their differentiation signals and inflammatory potential. In the presence of IL-1β and IL-23, or small intestinal serum amyloid A (SAA) proteins, IL-6-driven STAT3 activation promotes the polarization of pTh17 cells (11,12). These cells are highly inflammatory and induce severe disease in adoptive transfer models of experimental autoimmune encephalomyelitis (EAE), a mouse model of multiple sclerosis (11). In contrast, npTh17 cells polarized with IL-6 and TGF-β are less inflammatory. However, npTh17 cells can gain pathogenicity upon prolonged exposure to IL-23, further emphasizing the importance of IL-23 signaling in driving Th17-mediated autoimmune responses (10,13).

JunB, a member of the AP-1 transcription factor family, is a key regulator of Th17 cell differentiation and function. JunB expression is induced in response to IL-6 and is essential for the generation of IL-23-dependent pTh17 cells as well as the development of EAE and colitis (14,15). JunB regulates critical Th17 genes, including *Rorc*, *Il17a*, and *Il23r*, but its role in these genes is context-dependent: while JunB is necessary for *Il17a* expression in both pTh17 and npTh17 cells, its regulation of *Rorc* and *Il23r* is specific to pTh17 cells (14). Notably, JunB deficiency does not impair the development of gut-resident homeostatic Th17 cells, highlighting its specialized role in pathogenic Th17 subsets (14).

Because mature Th17 cells are the primary mediators of disease in autoimmune contexts, understanding the roles of transcription factors in these cells is essential. A recent study demonstrated that RORγt is essential for maintenance of mature Th17 cells by suppressing transdifferentiation into Th2-like cells (16). In contrast, roles of other transcription factors (TFs) critical for Th17 differentiation, such as JunB, remain unknown. To address this, we applied a targeted protein degradation system, the degradation tag (dTag) system, to acutely degrade JunB in mature Th17 cells. This approach provides important insights into JunB’s suitability as a therapeutic target and demonstrates the utility of the dTag system for dissecting the dynamic roles of transcription factors in immune cells.

## Methods

### Mice

Mice were housed under specific pathogen-free conditions at the Okinawa Institute of Science and Technology (OIST) animal center (Onna, Okinawa, Japan). All experimental procedures were approved by the Animal Care and Use Committee at OIST. All mice used were of the C57BL/6 background. OT-II (stock #004194) and B6.SJL (stock #002014) mouse strains were obtained from The Jackson Laboratory (Bar Harbor, ME, USA). FKBP12^F36V^-JunB (“dTAG-JunB”) and dTAG-JunB OT-II mice were generated in this study. Mice 6 to 12 weeks of age were used for all experiments.

### FKBP12^F36V^-JunB Mouse Engineering

FKBP12^F36V^ knock-in mice were generated by CRISPR-Cas9-mediated genome editing. A guide RNA (gRNA) targeting the *Junb* locus (sequence in Supplementary Table 1) was formed by duplexing crRNA with tracrRNA (Integrated DNA Technologies, Coralville, IA, USA). A single-stranded oligodeoxynucleotide (ssODN; sequence in Supplementary Table 1) was used as the homology-directed repair (HDR) template.

Embryos harvested from superovulated C57BL/6 females were electroporated with a mix of gRNA, Cas9 nuclease (Thermo Fisher Scientific, Waltham, MA, USA), and ssODN. Electroporation was performed using three 25 V pulses. After washing, embryos were cultured in potassium simplex optimized medium (KSOM) at 37°C, 5% CO[, and transferred into pseudo-pregnant ICR females. Pregnancy in recipient females was confirmed 14–15 days post-transfer.

### CD4+ Naïve T Cell Isolation

CD4+ naïve T cells were isolated from spleens of sacrificed mice. Spleens were mechanically dissociated, and red blood cells were lysed using ACK buffer (Thermo Fisher Scientific). CD4+ T cells were then enriched with the MojoSort™ Mouse CD4 T Cell Isolation Kit (BioLegend, San Diego, CA, USA) and stained with anti-CD3, anti-CD4, anti-CD44, and anti-CD62L (all antibodies from BioLegend). CD4+ CD25-CD62L^hi^ CD44^lo^ cells were sorted by flow cytometry using a FACSAria™ III (BD Biosciences, San Jose, CA, USA), excluding dead cells based on FSC/SSC gating. Sorted cells were resuspended in complete IMDM and kept on ice until use.

### Cell Culture

Naïve CD4+ T cells were plated at 0.5 x 10^6^ cells/mL in complete IMDM, consisting of IMDM (Invitrogen, Carlsbad, CA, USA) supplemented with 10% FBS (Sigma-Aldrich, St. Louis, MO, USA), 1x streptomycin-penicillin (Sigma-Aldrich), and 55 mM β-mercaptoethanol (Invitrogen). Cells were seeded on plates pre-coated with 5 μg/mL anti-CD3ε (clone 145-2C11; BioLegend) and cultured with 1 μg/mL anti-CD28 (clone 37.51; BioLegend).

To induce pTh17 cell polarization, IL-6 (20 ng/mL; BioLegend), IL-1β (20 ng/mL; BioLegend), and IL-23 (40 ng/mL; BioLegend) were added. For npTh17 cell polarization, IL-6 (20 ng/mL; BioLegend) and TGF-β (3 ng/mL; PeproTech, Cranbury, NJ, USA) were used. Cultures were incubated at 37°C with 5% CO_2_.

For cytokine expression analysis, cells were re-stimulated for 4–5 hours with phorbol 12-myristate 13-acetate (PMA; 50 ng/mL; Sigma-Aldrich) and ionomycin (500 ng/mL; Sigma-Aldrich), in the presence brefeldin A (5 μg/mL; BioLegend). For dTAG ligand treatment, cells were cultured with 400 nM dTAG^V^-1 (Bio-Techne, Minneapolis, MN, USA) or an equivalent volume of DMSO. Activin A supplementation was performed by adding 30 ng/mL activin A (BioLegend).

For experiments involving mature cells, 3-day polarized cells were transferred onto fresh, anti-CD3ε-coated plates and cultured in fresh complete IMDM containing anti-CD28 and the appropriate cytokines.

### Ovalbumin (OVA) Peptide Immunization

Naïve CD4+ T cells (0.5 x 10^6^ per mouse) isolated from dTAG-JunB OT-II mice (CD45.2+) were transferred into B6.SJL recipient mice (CD45.1+) by intravenous injection. The following day, each mouse was immunized subcutaneously at the base of the tail with 100 μg of OVA peptide 323-339 (GL Biochem, Shanghai, China) in 100 μL of a 1:1 PBS/CFA emulsion. CFA was prepared by suspending 100 mg of *Mycobacterium tuberculosis* H37Ra (BD Difco, Detroit, MI, USA) in 10 mL incomplete Freund’s adjuvant (BD Difco). The OVA/CFA emulsion was generated by vortexing a 1:1 mixture of antigen in PBS and CFA for 30 minutes at maximum speed.

### *Ex Vivo* Cell Culture

Seven days post-immunization, draining inguinal lymph nodes were collected and dissociated to obtain whole lymph node cell suspensions. Cells were cultured at 8 x 10^6^ cells/mL in complete IMDM containing 40 μg/mL OVA peptide 323-339 (GL Biochem). Cultures were treated with 400 nM dTAG ligand (Bio-Techne) or DMSO alone. For cytokine expression analysis, brefeldin A (5 μg/mL; BioLegend) was added during the final six hours of culture. Adoptively transferred dTAG-JunB OT-II CD4+ T cells were identified by flow cytometry as CD3+ CD4+ CD45.1-CD45.2+ cells.

### Flow Cytometry Analysis

Cells were stained on ice and in the dark for 20 minutes with Zombie NIR (BioLegend) to distinguish live from dead cells. For *ex vivo* samples, TruStain FcX™ (anti-mouse CD16/32, BioLegend) was included to block Fc receptor binding. After surface staining with fluorophore-conjugated antibodies, cells were fixed and permeabilized using the eBioscience™ Foxp3 / Transcription Factor Staining Buffer Set (Thermo Fisher Scientific) according to the manufacturer’s instructions. Intracellular staining was performed using the provided Permeabilization Buffer. Flow cytometry data were acquired on BD Aria II, Aria III, or Fortessa X-20 instruments (BD Biosciences) and analyzed using FlowJo software (FlowJo, LLC, Ashland, OR, USA). Live, single cells were gated using Zombie NIR staining and FSC/SSC parameters.

### Antibodies

For surface staining, antibodies against CD25 (clone PC61, BioLegend), CD45.2 (clone 104, BioLegend), and CD4 (clone GK1.5, BioLegend or BD Biosciences) were each used at a dilution of 1:400. Antibodies against CD45.1 (clone A20, BioLegend), CD3 (clone 17A2, BioLegend), CD44 (clone IM7, BioLegend), and CD62L (clone MEL-14, BioLegend) were also used at 1:400.

For intracellular staining, anti-JunB (clone C-11, Santa Cruz Biotechnology, Dallas, TX, USA) was used at a dilution of 1:200. An anti-mouse IgG1, κ antibody (BD Biosciences) was used as an isotype control for JunB staining in some experiments. Antibodies against IL-17A (clone TC11-18H10.1, BioLegend) and T-bet (clone 4B10, BioLegend) were used at 1:200, while antibodies against IFNγ (clone XMG1.2, BioLegend) and FoxP3 (clone 150D, BioLegend) were used at 1:400 and 1:100, respectively.

### RNA Isolation

Approximately 50,000 live cells were used for RNA isolation. If viability was below 85%, cells were stained with Zombie NIR (BioLegend) and sorted for live cells by flow cytometry. RNA was extracted using the RNAdvance Cell v2 kit (Beckman Coulter, Brea, CA, USA) following the manufacturer’s instructions. RNA concentration was measured with the Qubit™ RNA BR Assay Kit (Thermo Fisher Scientific) and stored at -80°C.

### Quantitative PCR (qPCR)

For qPCR, 100–300 ng of total RNA was reverse-transcribed using ReverTra Ace® qPCR RT Master Mix (Toyobo, Osaka, Japan) according to the manufacturer’s instructions. cDNA was diluted for use in qPCR reactions with KAPA SYBR® FAST qPCR Master Mix (Kapa Biosystems, Wilmington, MA). Reactions were run on a StepOnePlus Real-Time PCR System (Applied Biosystems, Foster City, CA, USA). Sequences of qPCR primers used are given in Supplementary Table 1. Ct values were normalized to *Actb*.

### Statistical Analyses

Data were plotted and analyzed using GraphPad Prism (GraphPad Software, San Diego, CA, USA). Bars represent mean ± SD. Statistical significance was determined using two-tailed Student’s t-tests or two-way ANOVA, with p-values indicated as ns (not significant), * (p<0.05), ** (p<0.01), *** (p<0.001), or **** (p<0.0001).

### RNA Sequencing & Differential Gene Expression Analysis

For RNA-seq, 10 ng of total RNA per sample was used to prepare libraries with the QuantSeq 3’ mRNA-Seq Library Prep Kit FWD (Lexogen, Vienna, Austria) following the manufacturer’s instructions. Libraries were sequenced (50 bp single-end reads) on Illumina platforms by the OIST Sequencing Section. Quality control of raw sequencing reads was performed using FastQC. Adapter trimming and quality filtering were conducted with Trimmomatic. Reads were aligned to the GRCm38 reference genome using HISAT2 and read counts were generated using featureCounts from the Subread package. Differentially expressed genes were identified using DESeq2.

### Induction of EAE

Mice were subcutaneously injected with 300 µg of myelin oligodendrocyte glycoprotein (MOG) 35–55 peptide (synthesized by PH Japan, Hiroshima, Japan) emulsified in 200 µL of CFA on day 0. CFA was prepared as described above. On days 0 and 2, mice were intraperitoneally injected with 400 ng of pertussis toxin (Funakoshi, Tokyo, Japan). Disease severity was assessed daily using the following scoring system: 1, limp tail; 2, limp tail and hind leg weakness; 3, limp tail with paralysis of one hind leg; 4, limp tail with paralysis of both hind legs; and 5, complete paralysis of both hind and front legs. Mice reaching a score of 5 were euthanized.

### Isolation of CNS-Infiltrating Mononuclear Cells

EAE-induced mice were sacrificed by CO[ asphyxiation and perfused intracardially with ice-cold PBS. Spinal cords and brains were isolated, minced, and digested in PBS containing 1 mg/mL collagenase D (Roche, Basel, Switzerland) and 2.5 mg/mL DNase I (Sigma-Aldrich) at 37°C for one hour with gentle shaking. The digested tissue was passed through a 100 μm cell strainer to obtain a single-cell suspension and subjected to density gradient centrifugation using 37% and 70% Percoll (GE Healthcare, Chicago, IL, USA). Mononuclear cells were collected from the interface, washed with PBS, incubated with Zombie NIR (BioLegend) and anti-Fc receptor blocking antibody (BioLegend), and then stained with antibodies against cell surface markers for flow cytometry analysis.

## Results

### dTAG Ligand Treatment of FKPB12^F36V^-JunB Cells Rapidly Induces JunB Degradation

To investigate the role of JunB in mature T cells, we decided to utilize the dTAG system. This targeted protein degradation system employs a mutant version of the FKBP12 protein, FKBP12^F36V^, expressed in-frame with the protein of interest (17). In the presence of a heterobifunctional molecule that binds both FKBP12^F36V^ and an E3 ubiquitin ligase complex, such as dTAG^V^-1, the protein of interest is targeted for polyubiquitination and subsequently degradation by the proteasome (18).

Using CRISPR-Cas9 genome editing, we designed a guide RNA (gRNA) to target a site upstream of the *Junb* translation start site. Alongside the Cas9-gRNA complex, a single-stranded oligodeoxynucleotide (ssODN) was introduced as a repair template for HDR. The ssODN contained homology arms flanking the Cas9-induced double-strand break (DSB), as well as sequences encoding a 2x hemagglutinin (HA)-tag, the FKBP12^F36V^ insert, and a linker sequence (Fig. 1A). Genotyping of the FKBP12^F36V^-JunB (“dTAG-JunB”) mice via PCR confirmed successful knock-in, yielding a 691 bp amplicon in homozygous animals (Fig. 1B). Flow cytometry analysis showed that JunB expression in *in vitro* polarized dTAG-JunB pathogenic Th17 (pTh17) cells was comparable, albeit slightly reduced, compared to wild type (WT) C57BL/6 cells (Fig. 1C). Knock-in also had no effect on the ratio of naïve CD4+ T cells found in both the spleen and in lymph nodes (Supplementary Fig. 1).

**Figure 1.**
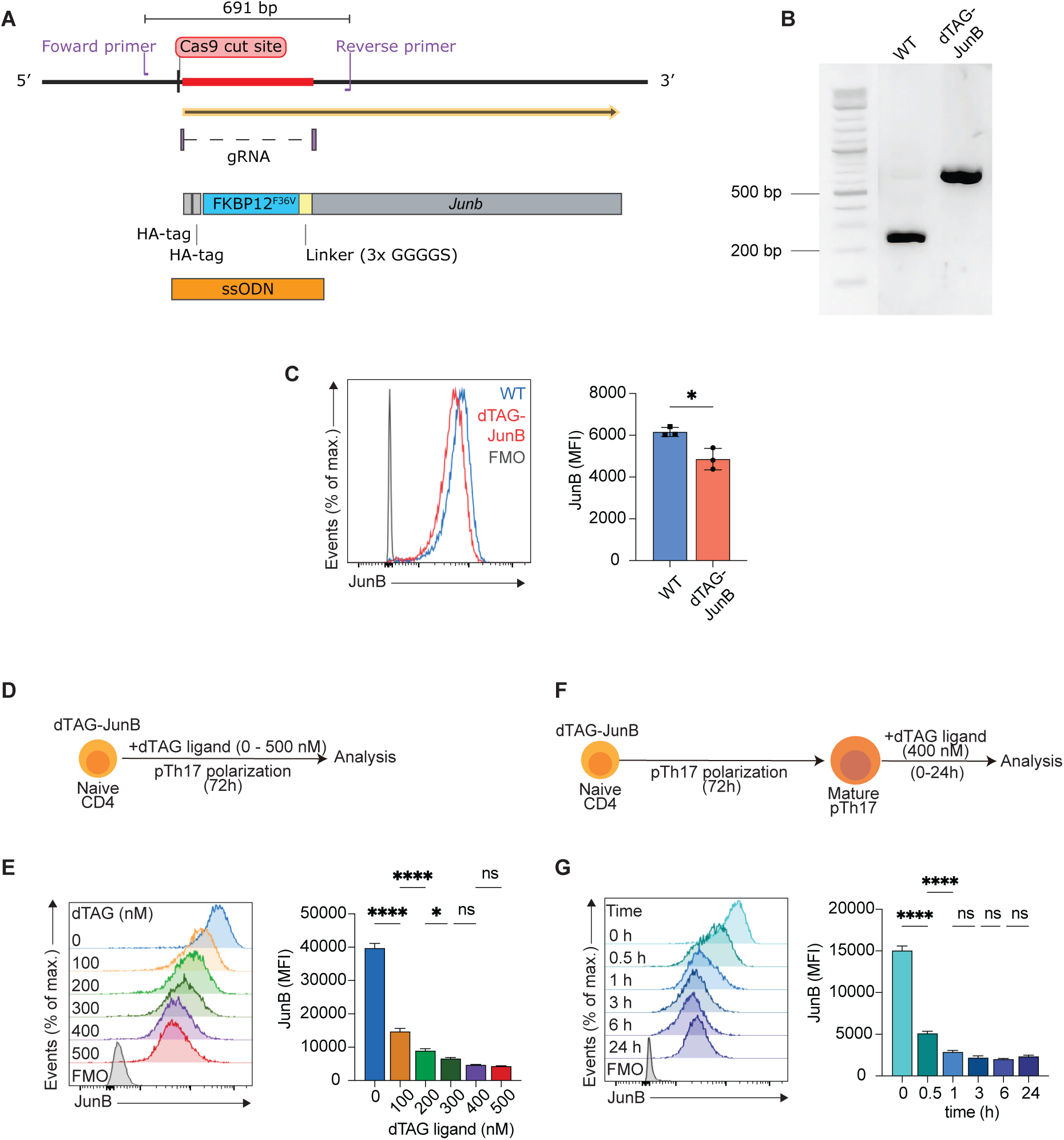
dTAG ligand induces rapid, dose-dependent degradation of JunB in FKBP12^F36V^ T cells *in vitro*. (**A**) Schematic of the *Junb* locus with the FKBP12^F36V^ insert (red line), enabling inducible JunB degradation by dTAG ligand. The insert was introduced via homology-directed repair (HDR) following a Cas9-mediated cut, using a single-stranded oligodeoxynucleotide (ssODN) template. gRNA = guide RNA; HA-tag = hemagglutinin tag. (**B**) Representative agarose gel showing PCR amplicons from WT and FKBP12^F36V^ (dTAG-JunB) mice. Primer annealing sites are indicated in (A). (**C**) JunB expression in WT and dTAG-JunB pTh17 cells polarized for 72 h, analyzed by flow cytometry. (**D**) Schematic of the experimental design for dTAG ligand treatment of dTAG-JunB cells during pTh17 polarization. (**E**) JunB expression in dTAG-JunB pTh17 cells treated as in (D), assessed by flow cytometry. (**F**) Schematic of the experimental design for dTAG ligand treatment of mature dTAG-JunB pTh17 cells. **(G)** JunB expression in mature dTAG-JunB pTh17 cells treated as in (F), analyzed by flow cytometry. Flow cytometry data are shown as mean ± SD from biological replicates, analyzed using two-tailed Student’s t-tests, and are representative of two independent experiments. P-values are indicated as ns (not significant), * (p<0.05), ** (p<0.01), *** (p<0.001), or **** (p<0.0001).

To confirm that dTAG^V^-1 (“dTAG ligand”) effectively degraded JunB in dTAG-JunB cells, we treated cells with increasing concentrations of the molecule during 72-hour polarization toward the pTh17 phenotype (Fig. 1D). JunB degradation occurred in a dose-dependent manner, with maximum degradation observed at 300 nM dTAG ligand (Fig. 1E). We then assessed the kinetics of degradation by adding dTAG ligand to 72-hour polarized pTh17 cells and measuring JunB expression over 24 hours (Fig. 1F). Within 30 minutes, JunB expression was significantly reduced, with maximal degradation achieved within one hour (Fig. 1G). These findings demonstrate that dTAG ligand treatment induces rapid and efficient degradation of JunB in dTAG-JunB T cells.

### dTAG Ligand Treatment Shows Minimal Side Effects

To confirm that the effects of dTAG ligand treatment were specific to JunB degradation, we treated WT pTh17 cells with dTAG ligand during polarization (Fig. 2A). As expected, dTAG ligand had no impact on JunB expression in WT cells (Fig. 2B), and treated cells displayed no changes in the expression of IL-17A or IFNγ (Fig. 2C).

**Figure 2.**
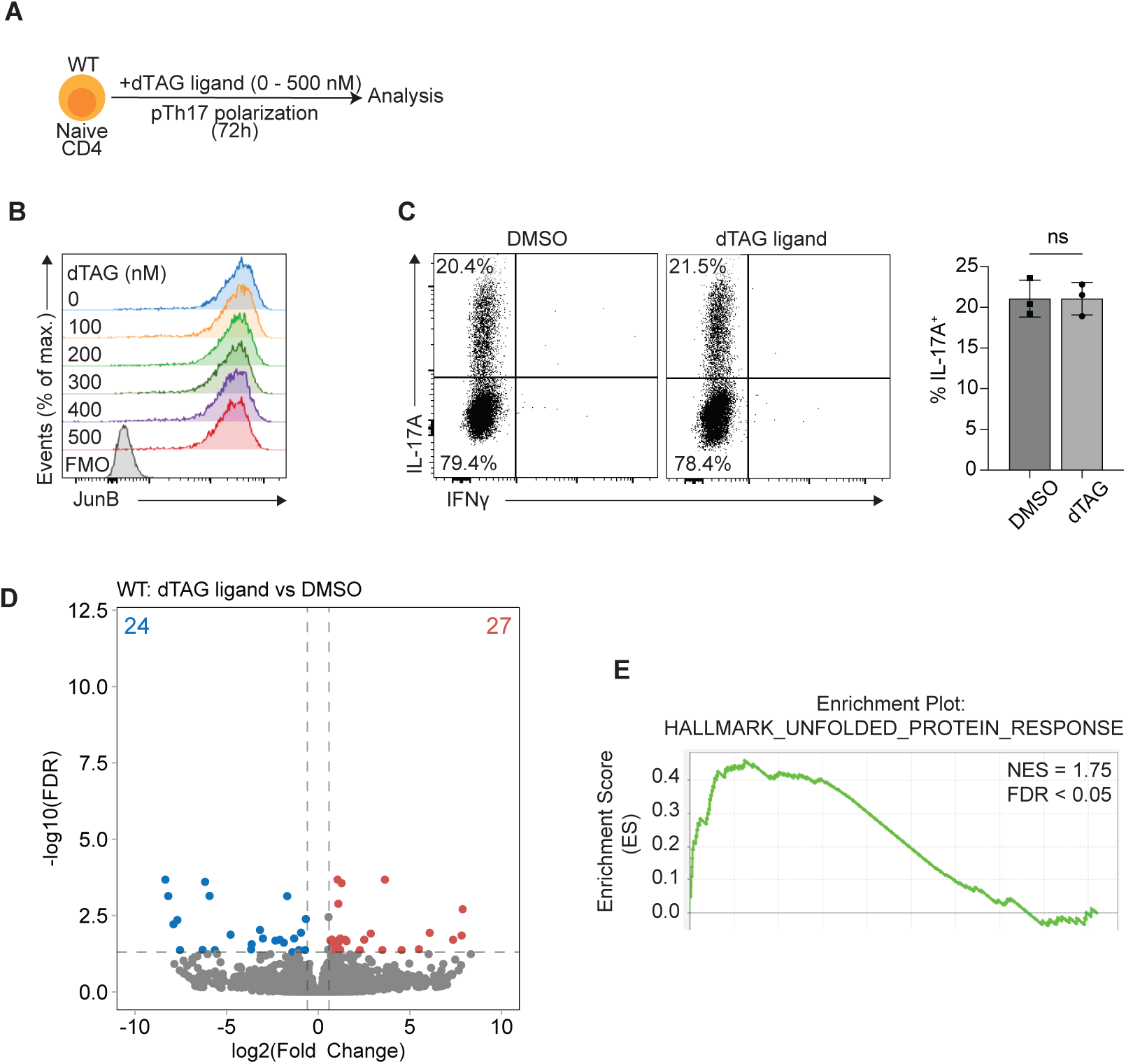
dTAG ligand has minimal off-target effects on WT Th17 cells. (**A**) Schematic of the experimental design for dTAG ligand treatment of WT cells during pTh17 polarization. (**B, C**) JunB expression **(B)** and cytokine expression (**C**) following re-stimulation in WT pTh17 cells treated as in (A). Data are shown as mean ± SD from biological replicates, as determined using the two-tailed Student’s t-test, and are representative of data from two independent experiments. P-values indicated as ns (not significant), * (p<0.05), ** (p<0.01), *** (p<0.001), or **** (p<0.0001). (**D**) Volcano plot of differentially expressed genes (DEGs) in WT pTh17 treated as in (A). Numbers of significantly down- and upregulated genes are shown in blue and red, respectively (FDR < 0.05, |log2 fold change| ≥ 0.59). Results are from 4 biological replicates. (**E**) Gene set enrichment analysis (GSEA) of the RNA-seq data described in (D). Normalized enrichment scores (NES) and FDR values are indicated.

To assess potential off-target effects at the transcriptomic level, we compared the global gene expression profiles of pTh17 cells treated with dTAG ligand or DMSO vehicle alone (Fig. 2D). A total of 51 differentially expressed genes (DEGs) were identified, none of which were key Th17-associated genes. Gene set enrichment analysis (GSEA) (19,20) of the RNA-seq data, using hallmark gene sets from the Molecular Signatures Database (21,22), revealed significant enrichment of only one gene set: the unfolded protein response (Fig. 2E). This finding suggests that dTAG ligand treatment may induce a mild endoplasmic reticulum stress response in pTh17 cells. Importantly, we observed no enrichment of gene sets associated with immune pathways, including the IFNγ response or IL-6/JAK/STAT3 signaling.

### dTAG-Mediated JunB Degradation Recapitulates Known Effects of JunB Deficiency in Polarizing pTh17 Cells

Previous studies have reported that *Cd4^Cre^Junb^fl/fl^* JunB-deficient (“JunB KO”) pTh17 cells exhibit altered expression of key Th17-associated molecules. Specifically, these cells show reduced IL-17A expression, increased IFNγ expression, elevated levels of the Th1 master regulator T-bet, and decreased expression of *Il17f*, *Rorc*, and *Il23r* (14,15). To determine whether dTAG ligand-mediated JunB degradation mimics these phenotypic changes, we analyzed dTAG-JunB cells treated with dTAG ligand during polarization toward the pTh17 phenotype (Fig. 3A).

**Figure 3.**
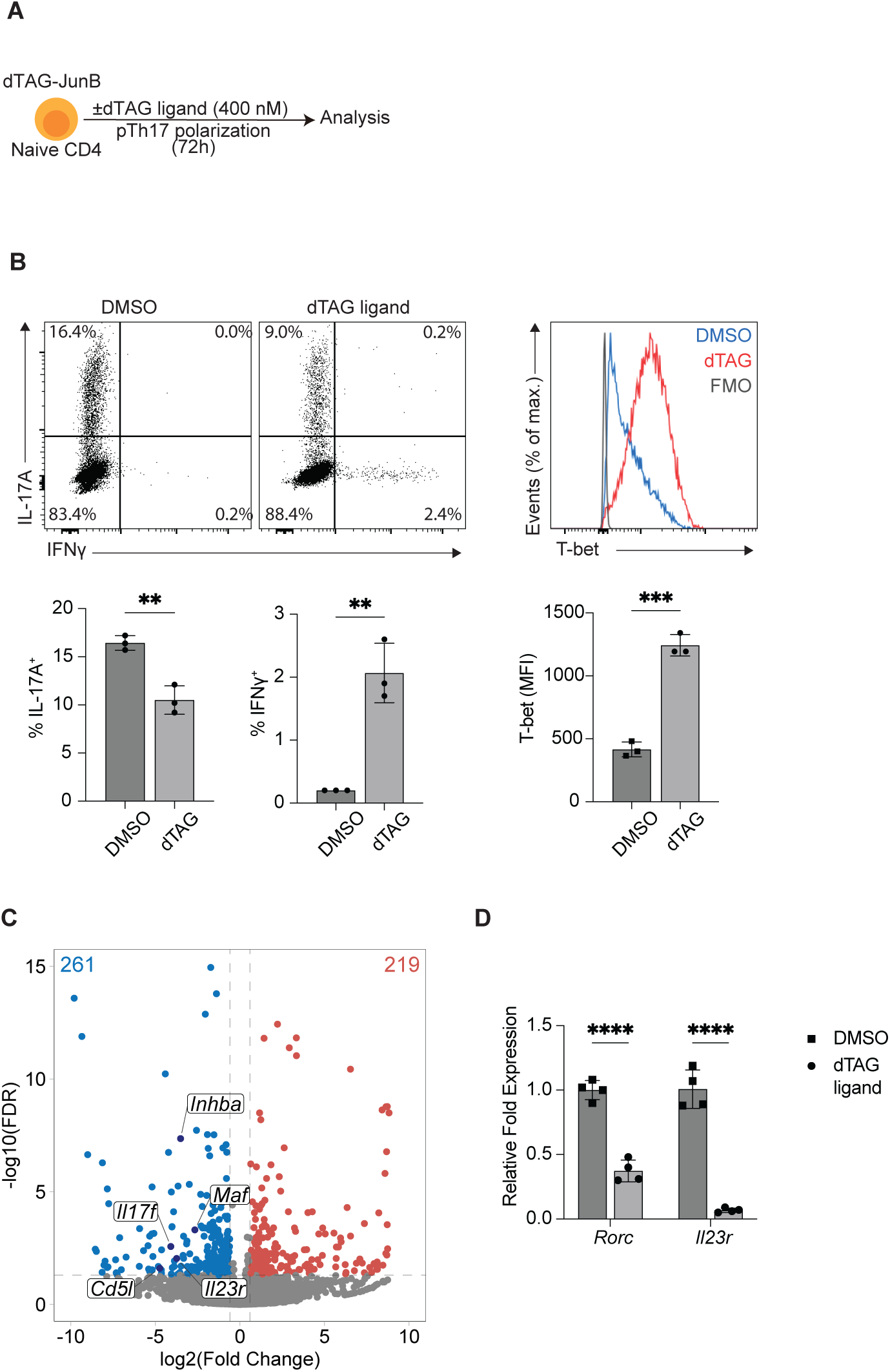
JunB degradation during pTh17 polarization reproduces known effects of JunB deficiency. (**A**) Schematic of the experimental design for dTAG ligand treatment of dTAG-JunB cells during pTh17 polarization. (**B**) Expression of cytokines (measured after a brief re-stimulation) and transcription factors in dTAG-JunB pTh17 cells treated as in (A), analyzed by flow cytometry. (**C**) Volcano plot of DEGs in dTAG-JunB pTh17 cells treated as in (A). Numbers of significantly down- and upregulated genes are shown in blue and red, respectively (FDR < 0.05, |log2 fold change| ≥ 0.59). Results are from 4 biological replicates. (**D**) mRNA expression of the indicated genes in dTAG-JunB pTh17 cells treated as in (A), assayed by qPCR and normalized to *Actb*. Flow cytometry and qPCR data are shown as mean ± SD from biological replicates, as determined using the two-tailed Student’s t-test, and are representative of data from two independent experiments. P-values indicated as ns (not significant), * (p<0.05), ** (p<0.01), *** (p<0.001), or **** (p<0.0001).

Consistent with findings in JunB KO cells, dTAG ligand-treated dTAG-JunB cells exhibited reduced IL-17A expression, increased IFNγ expression, and elevated T-bet levels (Fig. 3B). RNA-seq analysis further revealed decreased expression of *Il17f* and *Il23r* in dTAG ligand-treated cells (Fig. 3C). Although *Rorc* was not identified as a differentially expressed gene (DEG) in this analysis, qPCR confirmed a reduction in *Rorc* expression following dTAG ligand treatment (Fig. 3D). These findings demonstrate that dTAG-mediated JunB degradation effectively recapitulates the molecular changes associated with JunB deficiency in polarizing pTh17 cells.

### JunB is Required to Maintain the Phenotype of Mature Th17 Cells

To explore the role of JunB in mature Th17 cells, we cultured 72 h polarized pTh17 cells for an additional 48 h in the presence of dTAG ligand (Fig. 4A). Following JunB degradation, mature pTh17 cells exhibited increased T-bet expression and a higher percentage of IFNγ-positive cells after re-stimulation (Fig. 4B, 4C). These changes mirrored those observed in dTAG-JunB cells undergoing continuous dTAG ligand treatment during polarization (Fig. 3B). Interestingly, however, unlike during polarization, JunB degradation in mature pTh17 cells did not alter the percentage of IL-17A-expressing cells (Fig. 4B). This indicates that JunB is essential for IL-17A expression during differentiation but is dispensable for its maintenance in mature pTh17 cells. Using qPCR, we found that *Rorc* expression was unaffected by JunB degradation in mature pTh17 cells (Fig. 4D). This aligns with the established role of RORγt in sustaining IL-17A expression in mature Th17 cells (16). In contrast, *Il23r* expression was significantly reduced upon JunB degradation, highlighting its dependence on JunB in these cells.

**Figure 4.**
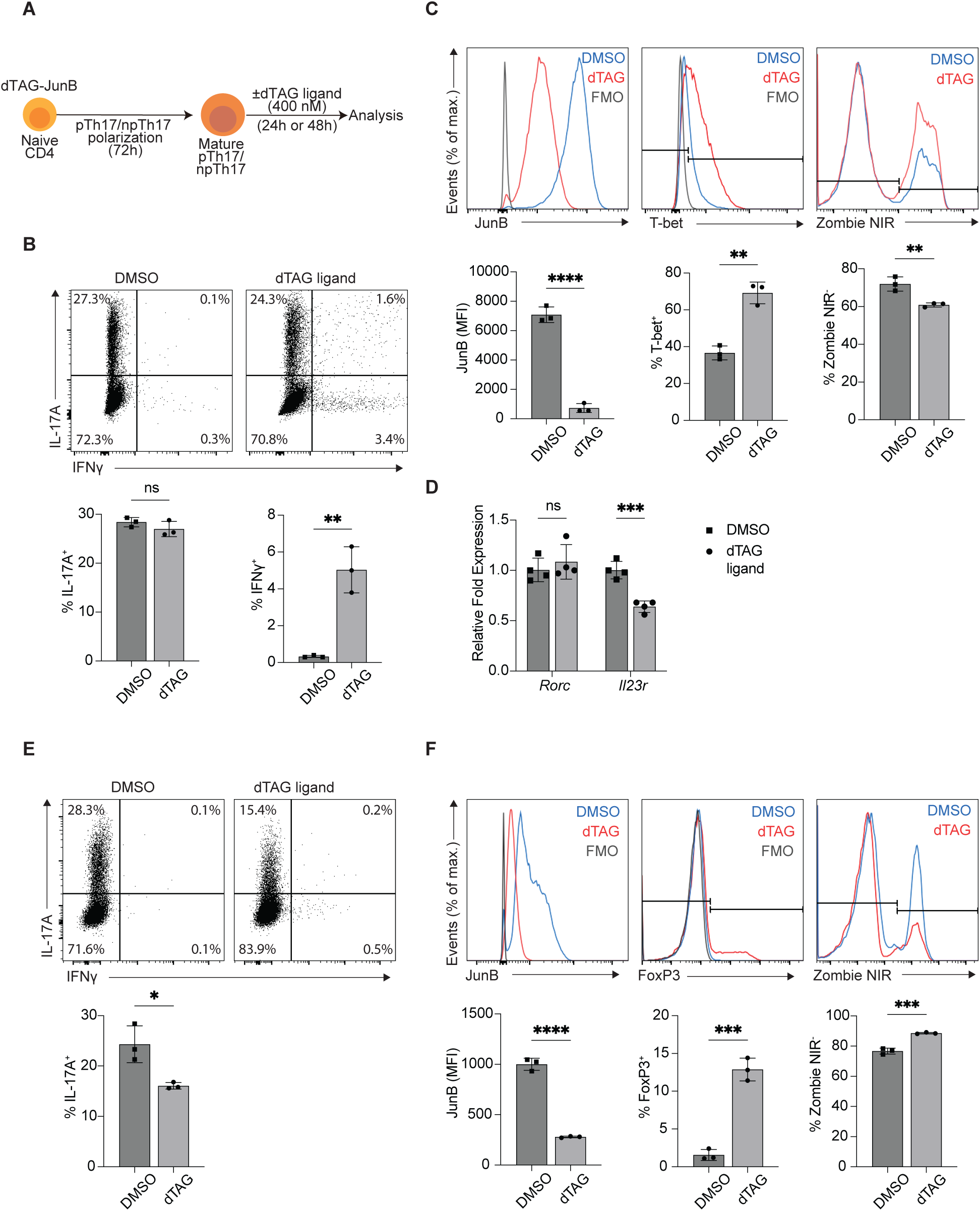
JunB degradation alters the phenotype of mature Th17 cells. (**A**) Schematic of the experimental design for dTAG ligand treatment of mature dTAG-JunB pTh17 cells. (**B, C**) Expression of cytokines (measured after a brief re-stimulation) **(B)** and transcription factors, and Zombie NIR dye staining (**C**) in dTAG-JunB pTh17 cells treated with dTAG ligand or DMSO for 48 h after an initial 72-hour polarization, analyzed by flow cytometry. (**D**) mRNA expression of the indicated genes in dTAG-JunB pTh17 cells treated with dTAG ligand or DMSO for 24 h after 72-hour polarization, assayed by qPCR and normalized to *Actb*. (**E, F**) Expression of cytokines (measured after a brief re-stimulation) **(E)** and transcription factors, and Zombie NIR dye staining (**F**) in dTAG-JunB npTh17 cells treated with dTAG ligand or DMSO for 48 h after an initial 72-hour polarization, analyzed by flow cytometry. Flow cytometry and qPCR data are shown as mean ± SD from biological replicates, as determined using the two-tailed Student’s t-test, and are representative of data from two independent experiments. P-values indicated as ns (not significant), * (p<0.05), ** (p<0.01), *** (p<0.001), or **** (p<0.0001).

Given a previous report that JunB supports the survival of Th cell subsets (23), we assessed the viability of mature pTh17 cells following JunB degradation. Loss of JunB resulted in a significant increase in cell death, as evidenced by enhanced Zombie NIR dye uptake (Fig. 4C), indicating that JunB promotes the viability of mature pTh17 cells.

In mature npTh17 cells, JunB degradation reduced IL-17A expression in these cells (Fig. 4E). Additionally, JunB degradation led to the induction of FoxP3, the Treg master regulator transcription factor, in a subset of cells (Fig. 4F). This role of JunB in repressing FoxP3 expression in npTh17 cells is consistent with its function during polarization, where JunB-deficient naïve cells polarized toward the npTh17 phenotype also exhibit elevated FoxP3 levels and reduced IL-17A expression (14,15). Unlike pTh17 cells, JunB degradation did not impair the viability of mature npTh17 cells. Instead, a slight increase in the percentage of Zombie NIR+ cells was observed (Fig. 4F), suggesting distinct roles for JunB in regulating cell death between these subsets.

These findings demonstrate that JunB is essential for maintaining the expression of key Th17 genes, including *Il23r*, in mature pTh17 cells, while being dispensable for others, despite its requirement during polarization. This underscores that JunB plays distinct roles in polarizing and mature pTh17 cells. Additionally, JunB degradation impairs pTh17 cell viability but not that of npTh17 cells.

### JunB Degradation in Mature Th17 Cells Induces Rapid Transcriptomic Changes

The rapidity of transcription factor degradation through the dTAG system allows for the identification of direct target effects before secondary responses or compensatory mechanisms arise (24,25). In dTAG-JunB cells, JunB degradation reaches its maximum within 1 hour of dTAG ligand treatment (Fig. 1G). To investigate the immediate effects of JunB depletion, we analyzed RNA from mature pTh17 cells treated with dTAG ligand for 6 hours or 24 hours (Fig. 5A).

**Figure 5.**
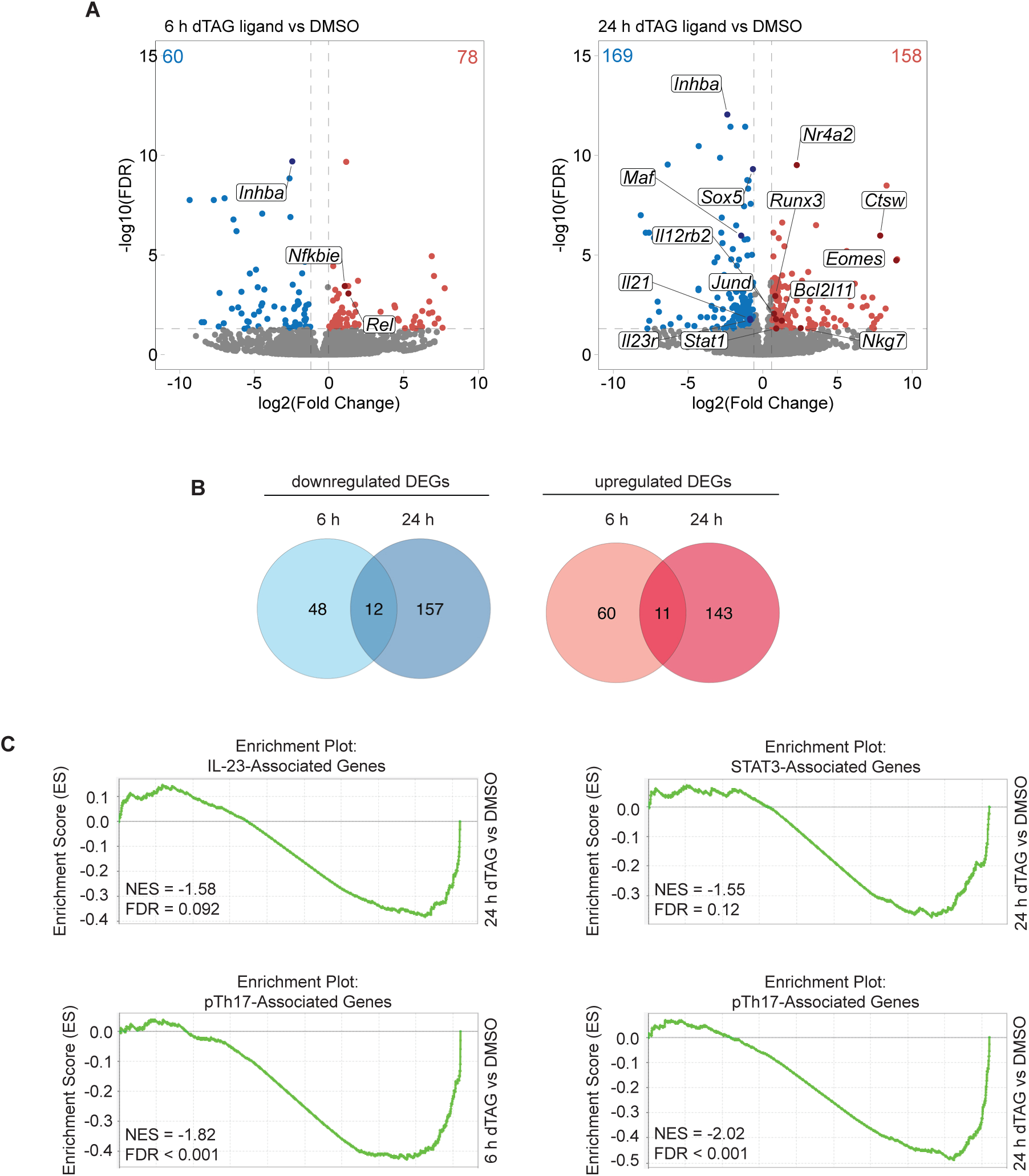
JunB degradation rapidly alters the mature pTh17 transcriptional program. (**A**) Volcano plots of DEGs in dTAG-JunB pTh17 cells treated with dTAG ligand or DMSO for 6 or 24 h after 72-hour polarization. Numbers of significantly down- and upregulated genes are shown in blue and red, respectively (FDR < 0.05, |log2 fold change| ≥ 0.59). Results are from 4 biological replicates. (**B**) Venn diagrams showing overlapping down- and upregulated DEGs identified in (A). (**C**) GSEA of RNA-seq data described in (A). NES and FDR values are indicated.

After 6 hours of treatment, RNA-seq analysis revealed 60 downregulated and 78 upregulated DEGs, indicating rapid transcriptomic changes. Genes associated with key Th17 signaling pathways were affected, including upregulation of *Rel* and *Nfkbie* (involved in NF-κB signaling) and downregulation of *Inhba*, which encodes the βA subunit of activin A. After 24 hours, the number of DEGs increased to 169 downregulated and 158 upregulated genes. Notably, only a subset of DEGs identified at 6 hours persisted at 24 hours, suggesting transient gene expression changes (Fig. 5B). Persistent downregulated genes included *Inhba*, *Il33*, and *Il2ra*, while *Il2rn*, *Ccl4*, and *Gbp2* remained upregulated. The pro-apoptotic gene *Bcl2l11* was also upregulated, consistent with JunB’s role in maintaining pTh17 cell viability (Fig. 4C).

JunB has been implicated in restraining alternative T helper fates, particularly the Th1 phenotype during inflammation (15). JunB degradation in mature pTh17 cells upregulated several Th1-associated genes, including *Il12rb2*, *Stat1*, and *Runx3* (26,27). Additionally, genes linked to Th1-like ex-Th17 cells, such as *Eomes*, *Ctsw*, and *Nkg7* (28,29), were strongly upregulated. *Eomes*, a key regulator of Th17-to-Th1 conversion, was among the most significantly upregulated genes after 24 hours of dTAG ligand treatment.

To further understand these changes, we performed GSEA using Th17 gene sets defined by Lee et al. (30). JunB degradation led to the negative enrichment of genes downregulated in the absence of IL-23 signaling (Fig. 5C), consistent with reduced *Il23r* expression. This suggests that JunB loss shifts the Th17 transcriptome toward IL-1β and IL-6-polarized states with diminished IL-23 influence. Additionally, a gene set associated with pTh17 polarization conditions showed reduced enrichment following JunB degradation, indicating a loss of pTh17 cell identity. Remarkably, this shift toward a more Th0-like profile occurred even after only six hours of treatment. Further GSEA using a STAT3-dependent gene set associated with IL-23 signaling (7) revealed that the transcriptomic profile of pTh17 cells following JunB degradation resembled that of STAT3-deficient cells.

RNA-seq analysis in mature npTh17 cells revealed a rapid downregulation of the Th17 cytokines *Il17a* and *Il17f* within 6 hours of JunB degradation (Supplementary Fig. 2A). Interestingly, *Il9* expression was upregulated by 24 hours. IL-9 is known to synergize with TGF-β to promote the differentiation of Th17 cells from naïve CD4+ T cells, and IL-9 receptor-deficient mice exhibit less severe EAE symptoms (18). Similar to observations in pTh17 cells, only a subset of DEGs identified at 6 hours persisted at 24 hours, indicating that many gene expression changes induced by JunB degradation are transient (Supplementary Fig. 2B).

These results demonstrate that rapid JunB degradation alters the transcriptional landscape of mature pTh17 and npTh17 cells, shifting pTh17 cells away from an IL-23–dependent Th17 identity and inducing transient gene expression changes in both subsets.

### JunB Promotes *Il23r* Expression in *In vivo* Activated Th17 Cells

Having observed the effects of JunB degradation in mature pTh17 cells under *in vitro* polarization conditions, we next investigated whether JunB played a similar role in Th17 cells activated *in vivo*. Although dTAG^V^-1has been successfully administered to mice to induce target protein degradation *in vivo* (18), our attempts to degrade JunB in CNS-infiltrating CD4+ T cells of EAE-induced dTAG-JunB mice were unsuccessful (Supplementary Fig. 3). We therefore employed an *ex vivo* approach.

To this end, we generated dTAG-JunB OT-II transgenic mice by crossing dTAG-JunB mice with OT-II mice, which express a TCR specific for the I-Ab-restricted chicken OVA peptide 323-339. Naïve CD4+ T cells (CD45.2+) from dTAG-JunB OT-II mice were transferred into congenic B6.SJL recipient mice (CD45.1+) via intravenous injection. The following day, recipient mice were immunized with OVA peptide emulsified in CFA, which is known to predominantly induce a Th17 response (23). Seven days post-immunization, draining inguinal lymph nodes were harvested, and cell suspensions were cultured *ex vivo* in the presence of OVA peptide, along with either dTAG ligand or DMSO alone (Fig. 6A).

**Figure 6.**
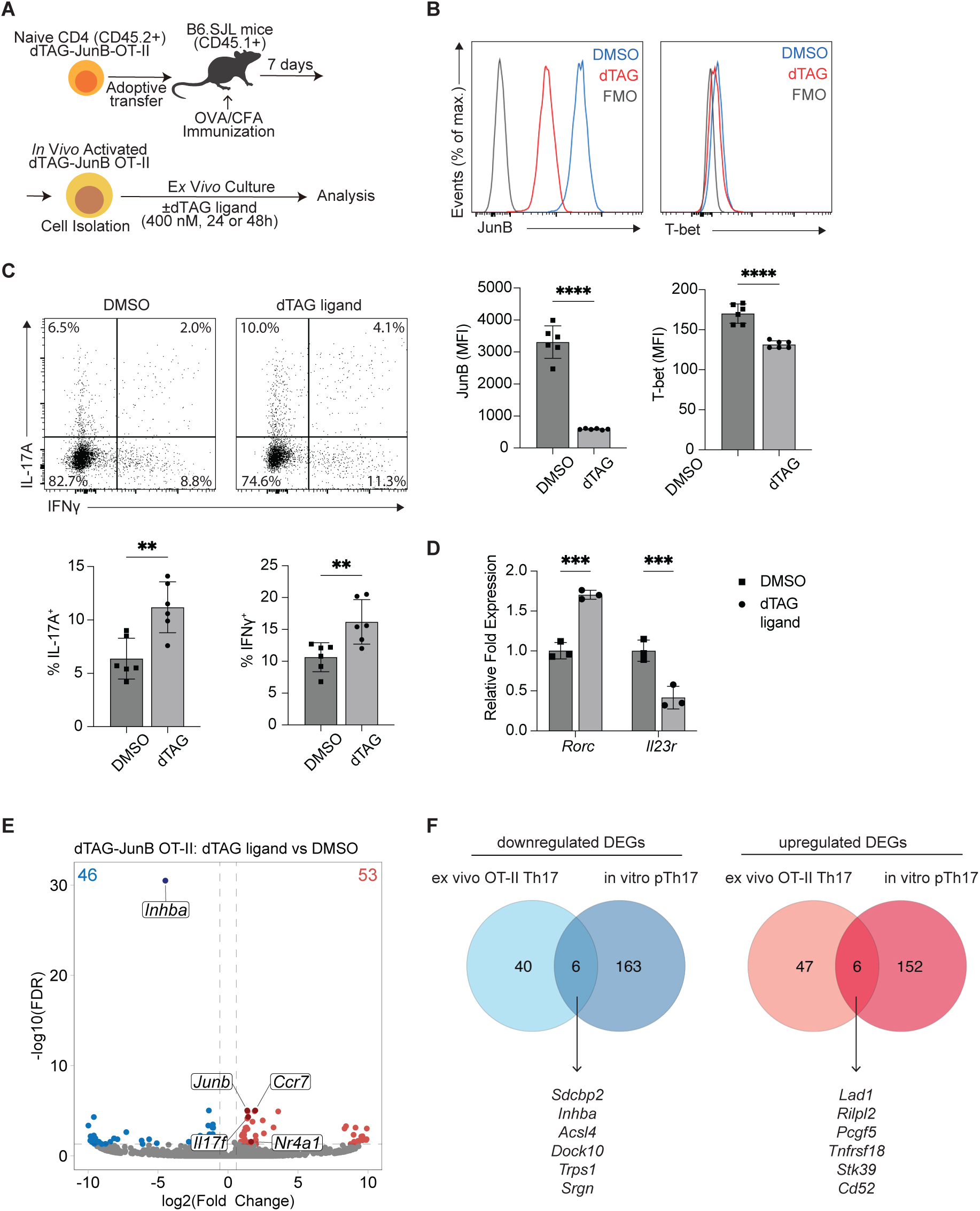
*Ex vivo* dTAG ligand treatment of *in vivo* activated dTAG-JunB Th17 cells alters expression of key Th17 molecules. (**A**) Schematic of the experimental design for *ex vivo* dTAG ligand treatment of dTAG-JunB OT-II cells isolated from draining lymph nodes of recipient mice after adoptive transfer and OVA/CFA immunization. (**B, C**) Expression of transcription factors (**B**) and cytokines (measured after a brief re-stimulation) (**C**) in dTAG-JunB OT-II cells following *ex vivo* treatment with dTAG ligand or DMSO for 48 h. (**D**) mRNA expression of the indicated genes in isolated dTAG-JunB OT-II cells cultured *ex vivo* with dTAG ligand or DMSO for 24 h, assayed by qPCR and normalized to *Actb*. Flow cytometry and qPCR data are shown as mean ± SD from biological replicates, as determined using the two-tailed Student’s t-test, and are representative of data from two independent experiments. P-values indicated as ns (not significant), * (p<0.05), ** (p<0.01), *** (p<0.001), or **** (p<0.0001). (**E**) Volcano plot of DEGs in dTAG-JunB OT-II cells treated as in (D). Numbers of significantly down- and upregulated genes are shown in blue and red, respectively (FDR < 0.05, |log2 fold change| ≥ 0.59). Results are from 3 biological replicates. (**F**) Venn diagrams showing overlapping down- and upregulated DEGs following dTAG ligand treatment of mature *in vitro* polarized dTAG-JunB pTh17 cells (from Fig. 5A, 24 h treatment) and *ex vivo* treated dTAG-JunB OT-II cells (from Fig. 6E).

We confirmed that dTAG-JunB OT-II cells expressed high levels of JunB, which was effectively degraded following *ex vivo* dTAG ligand treatment (Fig. 6B). Similar to our findings in *in vitro* polarized pTh17 cells, JunB was not required for IL-17A expression in these *in vivo* activated cells (Fig. 6C). Interestingly, JunB degradation increased the number of IL-17A+ cells. This increase may be explained by elevated *Rorc* expression observed upon JunB degradation (Fig. 6D). In contrast, JunB degradation resulted in a higher number of IFNγ+ cells (Fig. 6C) and downregulated *Il23r* (Fig. 6D), a finding consistent with the results seen in *in vitro* polarized pTh17 cells.

Finally, transcriptomic analysis of *ex vivo* cultured dTAG-JunB OT-II cells identified 99 DEGs after 24 hours of dTAG ligand treatment (Fig. 6E), with 12 overlapping DEGs shared with *in vitro* polarized pTh17 cells (Fig. 6F). Among these, *Inhba* was consistently downregulated, underscoring JunB’s importance in activin A signaling across both *in vitro* and *in vivo*-activated Th17 cell contexts.

These findings demonstrate that JunB also promotes *Il23r* expression in Th17 cells activated *in vivo*, reflecting its role under *in vitro* conditions. Furthermore, the consistent downregulation of *Inhba* in both contexts highlights JunB’s involvement in regulating activin A signaling in Th17 cells.

### JunB is Dispensable for pTh17 Cell Polarization in the Presence of Activin A

*Inhba* is induced during CD4+ T cell polarization by a cytokine cocktail of IL-6, IL-1β, and IL-23 and has been shown to enhance Th17 pathogenicity (30). Suppressing activin A signaling during Th17 polarization reduces the expression of key Th17 genes, including *Rorc*, *Il17a*, and *Il23r*. Given that JunB degradation leads to reduced *Inhba* expression, we hypothesized that JunB-mediated upregulation of *Inhba* is critical for pTh17 cell polarization.

To test whether activin A supplementation could rescue the effects of JunB degradation, we added activin A to the cell culture medium during pTh17 polarization (Fig. 7A). With activin A supplementation, the percentage of IL-17A+ cells was unaffected by JunB degradation (Fig. 7B). Interestingly, activin A supplementation increased the percentage of IL-17A+ cells even in control conditions. However, JunB degradation still resulted in an increased percentage of IFNγ+ cells, regardless of activin A supplementation. Additionally, while activin A partially reversed the increase in T-bet+ cells induced by JunB degradation (Fig. 7C), it restored *Rorc* expression but had no effect on *Il23r* expression (Fig. 7D). In contrast, in mature pTh17 cells treated with activin A (Fig. 7E), the percentage of IL-17A+ cells was not affected (Fig. 7F), whereas the percentage of T-bet+ cells induced by JunB degradation decreased (Fig. 7G). Furthermore, the expression levels of *Rorc* and *Il23r* were unaffected (Fig. 7H), indicating that activin A does not alter expression of these genes in mature cells.

**Figure 7.**
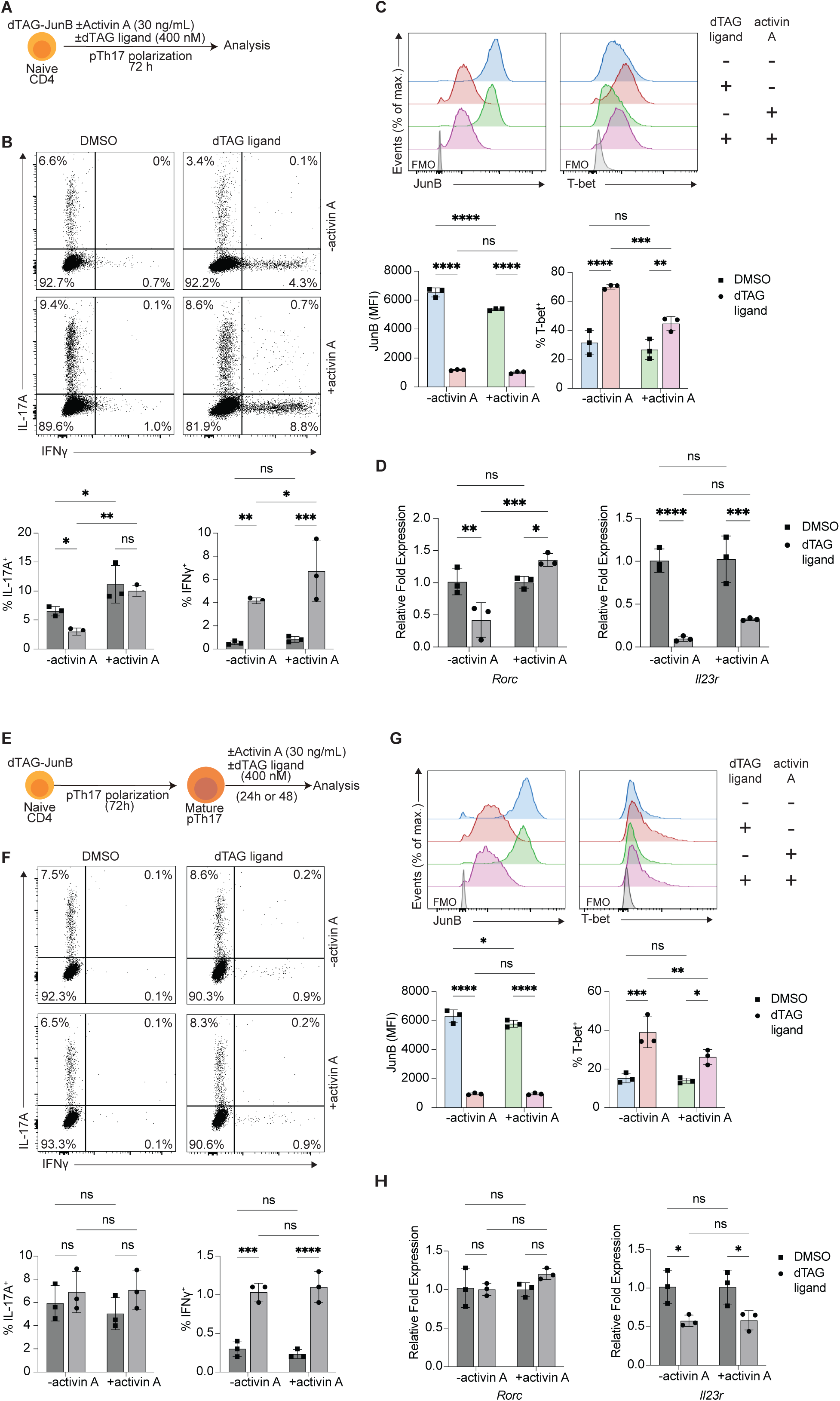
Activin A supplementation rescues effects of JunB loss during pTh17 polarization. (**A**) Schematic of the experimental design for dTAG ligand and/or activin A treatment of dTAG-JunB cells during pTh17 polarization. (**B, C**) Expression of cytokines (measured after a brief re-stimulation) **(B)** and transcription factors (**C**) in dTAG-JunB pTh17 cells polarized for 72 h and treated as in (A), analyzed by flow cytometry. (**D**) mRNA expression of the indicated genes in dTAG-JunB pTh17 cells treated as in (A), assayed by qPCR and normalized to *Actb*. (**E**) Schematic of the experimental design for dTAG ligand and/or activin A treatment of mature dTAG-JunB pTh17 cells. (**F, G**) Expression of cytokines (measured after a brief re-stimulation) **(F)** and transcription factors (**G**) in dTAG-JunB pTh17 cells polarized for 72 h and then cultured an additional 48 h with or without dTAG ligand and/or activin A, analyzed by flow cytometry. (**H**) mRNA expression of the indicated genes in dTAG-JunB pTh17 cells polarized for 72 h and then cultured an additional 24 h with or without dTAG ligand and/or activin A, as assayed by qPCR and normalized to *Actb*. Flow cytometry and qPCR data are shown as mean ± SD from biological replicates, as determined using two-way ANOVA, and are representative of data from two independent experiments. P-values indicated as ns (not significant), * (p<0.05), ** (p<0.01), *** (p<0.001), or **** (p<0.0001).

Together, these data indicate that JunB-mediated activin A expression is critical for driving the pTh17 phenotype during polarization by upregulating *Il17a* and *Rorc*. However, activin A may be dispensable for maintaining these genes in mature pTh17 cells.

## Discussion

Th17 cells are key mediators of autoimmune diseases, with their inflammatory potential critically dependent on IL-23 signaling. Understanding the transcriptional control of Th17 cells is important for developing targeted therapies for conditions such as multiple sclerosis, IBD, and psoriasis. JunB has been established as crucial for pTh17 cell generation, with JunB-deficient mice exhibiting resistance to Th17-mediated autoimmune diseases. However, studies utilizing T cell-specific *Junb* KO models have been unable to address whether JunB is required to maintain the phenotype of mature Th17 cells.

This study demonstrates the application of the dTAG system to degrade endogenous protein in T cells, focusing on JunB. Using WT Th17 cells, we demonstrated that the dTAG ligand molecule has minimal indirect effects, validating its use for studying T cell proteins. The dTAG system enables rapid, ligand-induced protein degradation, providing precise temporal control to explore the immediate effects of protein loss. By applying dTAG-mediated JunB degradation during pTh17 polarization, we confirmed JunB’s established role in regulating the pTh17 cell phenotype. Extending this analysis to mature T cells, we found that JunB is critical for maintaining phenotypic stability of pTh17 cells. In mature pTh17 cells, JunB degradation induced a shift toward a more Th1-like phenotype, which has been associated with pathogenicity in autoimmune diseases (31). Conversely, in mature npTh17 cells, JunB degradation resulted in a shift toward a FoxP3+ and IL-9+ phenotype, which may confer protective effects in autoimmune contexts (32).

JunB regulates Th17 phenotypes through distinct mechanisms depending on the differentiation state of the cells. During pTh17 polarization, JunB-mediated upregulation of *Inhba* (encoding activin A) was critical for sustaining *Rorc* expression. However, in mature pTh17 cells, JunB was dispensable for maintaining *Rorc*. Interestingly, JunB was not required for IL-17A expression in mature pTh17 cells but was necessary in mature npTh17 cells, possibly due to the induction of FoxP3, a known inhibitor of RORγt (33). Importantly, JunB was essential for *Il23r* expression in mature pTh17 cells, and its degradation led to reduced *Il23r* expression and diminished enrichment of an IL-23 signaling-associated gene set, as shown by GSEA. This function appears independent of JunB’s role in promoting activin A signaling.

Beyond IL-23 signaling, JunB influences Th17 cell survival, with distinct effects in pTh17 and npTh17 subsets. In mature pTh17 cells, JunB degradation reduced cell viability, correlating with increased expression of the pro-apoptotic gene *Bcl2l11*, as revealed by RNA-seq analysis (Fig. 5A). A previous study has shown that JunB represses *Bcl2l11* by promoting IRF4 binding to its regulatory regions (23). Conversely, in mature npTh17 cells, JunB degradation slightly enhanced viability, accompanied by upregulation of *Il9*, a cytokine known to protect T cells from apoptosis through STAT3- and STAT5-dependent mechanisms (34). These findings indicate that JunB’s role in regulating Th17 cell survival is context-dependent, differing between pathogenic and non-pathogenic subsets.

We were unable to observe degradation of JunB in CNS-infiltrating CD4+ T cells following administration of dTAG ligand to mice with induced EAE. This may be due to the impermeability of the blood-brain barrier (BBB) to the dTAG ligand molecule (35), although the BBB does become disrupted and more permeable during EAE (36). Alternatively, the dose, timing and/or route of dTAG ligand administration used here may have been insufficient to achieve JunB degradation. Future work could optimize dTAG ligand administration for *in vivo* studies and validate JunB’s therapeutic potential by assessing its role in ongoing Th17-mediated disease models, such as EAE.

Collectively, these results position JunB as a potential therapeutic target for Th17-driven diseases. By sustaining *Il23r* expression, JunB promotes IL-23 responsiveness, a critical factor in Th17-mediated autoimmune responses. Moreover, targeting JunB could reduce the viability of inflammatory Th17 cells while blocking JunB-driven activin A secretion, which may exacerbate inflammation by promoting the polarization of neighboring inflammatory Th17 cells. This study also underscores the utility of the dTAG system as a powerful tool for dissecting transcription factor dynamics in T cells. Future applications of this system hold promise for paving the way towards innovative therapies targeting transcriptional regulators in autoimmune diseases.

## Acknowledgements

We are grateful for the assistance and support provided by the Core Facilities at OIST. We thank the Animal Resources Section for their support with mouse housing and care, the Instrument Analysis Section for providing access to and training on the flow cytometer instruments, the OIST Sequencing Section for their assistance with sample sequencing, and the Scientific Computing and Data Analysis Section for providing computing resources.

