## Supplementary Figures and Tables for "Delineating JunB’s Crucial Function in Mature Th17 Cells through Inducible Targeted Protein Degradation"

### Supplementary Figure 1

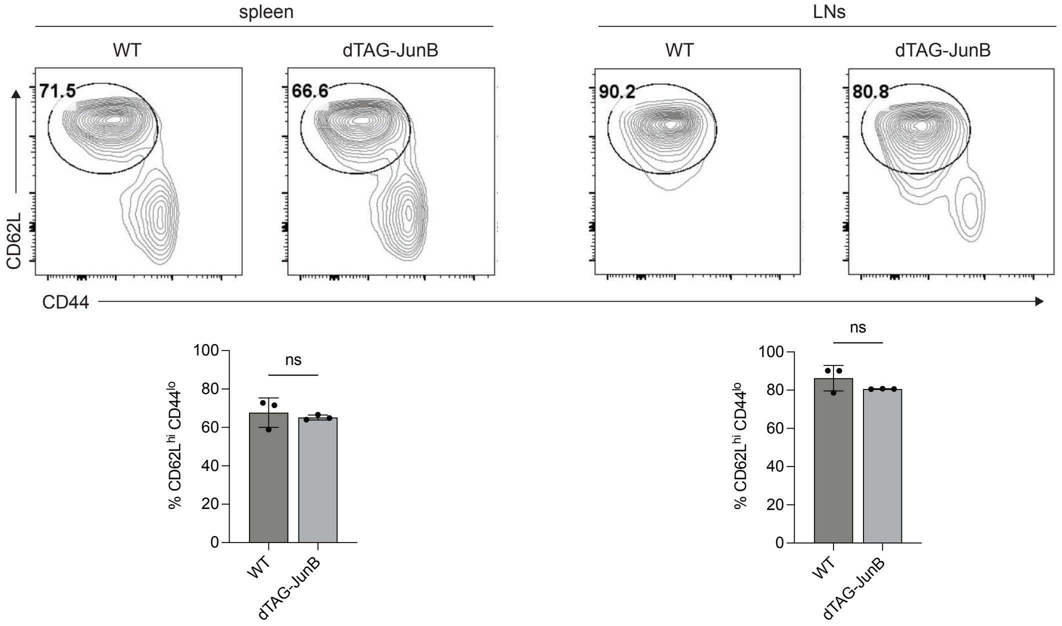


**Supplementary Figure 1. FKBP12$^{F36V}$ knock-in at the Junb locus does not affect naïve CD4+ T cell ratios.**

CD62L and CD44 staining of CD4+ T cells isolated from the spleen or pooled lymph nodes (LNs) of WT or dTAG-JunB mice, analyzed by flow cytometry. Data are presented as mean ± SD from biological replicates. Statistical significance was determined using a two-tailed Student's t-test. p-values are indicated as ns (not significant), * ($p<0.05$), ** ($p<0.01$), *** ($p<0.001$), **** ($p<0.0001$).

### Supplementary Figure 2

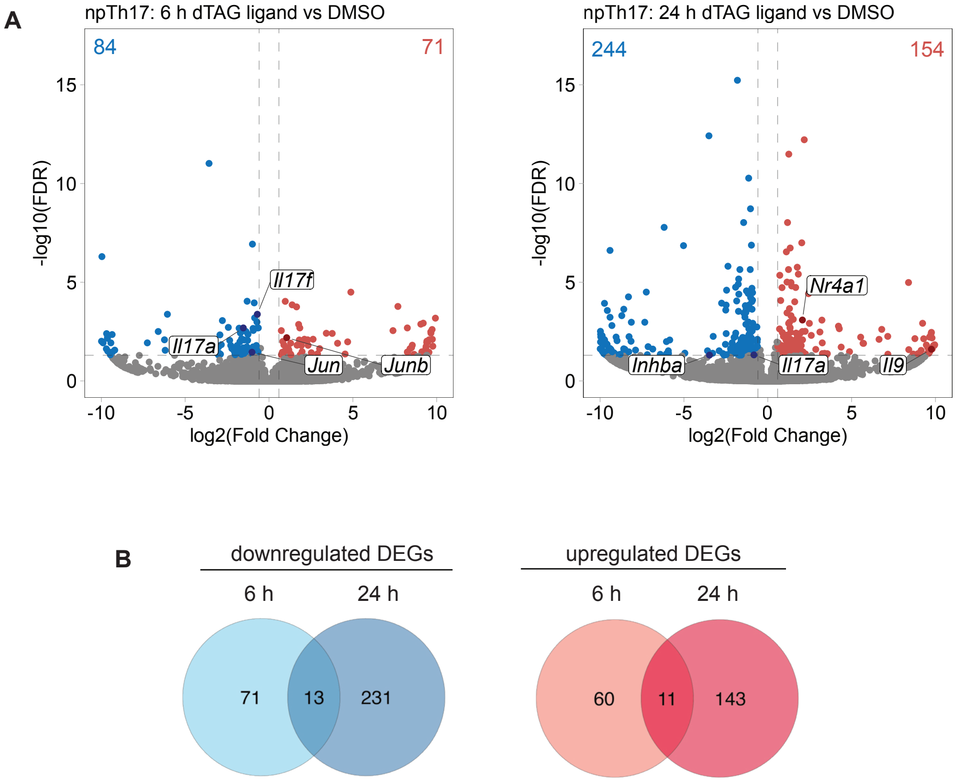


**Supplementary Figure 2. JunB degradation rapidly alters the mature npTh17 transcriptional program.**
(**A**) Volcano plots showing differentially expressed genes (DEGs) in dTAG-JunB npTh17 cells treated with dTAG-$^{V}$1 or DMSO for 6 or 24 hours following 72-hour polarization. Numbers of significantly down- and upregulated genes are shown in blue and red, respectively (FDR < 0.05, |log2 fold change| ≥ 0.59). Results are from 3 biological replicates. (**B**) Venn diagrams illustrating overlapping down- and upregulated DEGs identified in (A).

### Supplementary Figure 3

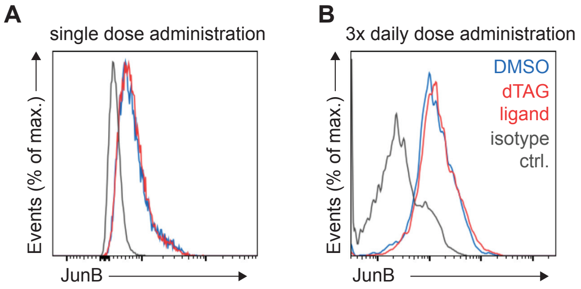


**Supplementary Figure 3. *In vivo* dTAG ligand administration fails to degrade JunB in CNS-infiltrating dTAG-JunB CD4+ T cells.**

JunB staining of CNS-infiltrating CD4+ T cells isolated from EAE-induced dTAG-JunB mice. Approximately two weeks after MOG 35–55 immunization, mice received one dose (A) or three daily doses (B) of 35 mg/kg dTAG ligand or DMSO vehicle intraperitoneally. In (A), mice were sacrificed 5 hours after the single dose; in (B), mice were sacrificed 24 hours after the final dose. Mononuclear cells were isolated from the CNS and analyzed by flow cytometry. An aliquot of cells was stained with a fluorophore-conjugated isotype IgG antibody as a control. The data show gated viable, CD3+ CD4+ cells.

**Supplementary Table 1. List of oligos used in this study.**

| Oligo | Sequence |
|---|---|
| gRNA for CRISPR/Cas9-mediated DSB at the *Junb* locus | CCCGGATGTGCACGAAAATG |
| ssODN for $FKBP12^{F36V}$ knock-in at the *Junb* locus | AGGGGGACCTTTCCCAGATCGCCCAGGCCGCCCGGATGTACCCCTAC GACGTGCCCGACTACGCCGGCTATCCGTATGATGTCCCGGACTATGC AGGAAGCGGAGGAGTGCAGGTGGAAACCATCTCCCCAGGAGACGGGC GCACCTTCCCCAAGCGCGGCCAGACCTGCGTGGTGCACTACACCGGG ATGCTTGAAGATGGAAAGAAAGTTGATTCCTCCCGGGACAGAAACAA GCCCTTTAAGTTTATGCTAGGCAAGCAGGAGGTGATCCGAGGCTGGG AAGAAGGGGTTGCCCAGATGAGTGTGGGTCAGAGAGCCAAACTGACT ATATCTCCAGATTATGCCTATGGTGCCACTGGGCACCCAGGCATCAT CCCACCACATGCCACTCTCGTCTTCGATGTGGAGCTTCTAAAACTGG AAGGTGGCGGTGGCTCGGGCGGTGGTGGGTCGGGTGGCGGCGGATCT TGCACGAAAATGGAACAGCCTTTCTATCACGACGACTCT |
| mouse genotyping: dTAG-JunB forward primer | TCTCCAGCTCCCGAGGACG |
| mouse genotyping: dTAG-JunB reverse primer | CAGGTTGAGCGCCAAGGTG |
| qPCR: *Il23r* forward primer | GTCCACCAAACTTCCCAGACAG |
| qPCR: *Il23r* reverse primer | CCTGAAGCAGGATGTCCTCTGA |
| qPCR: *Rorc* forward primer | AGCGCACCAACCTCTTTTCAC |
| qPCR: *Rorc* reverse primer | ATGAAGCCTGAAAGCCGCTTG |
| qPCR: *Actb* forward primer | CATTGCTGACAGGATGCAGAAGG |
| qPCR: *Actb* reverse primer | TGCTGGAAGGTGGACAGTGAGG |
